# Cross-disease comparison of dermatomyositis and lupus skin identifies inflammatory monocytes and JAK-1 signaling as drivers of vasculopathy in dermatomyositis

**DOI:** 10.1101/2025.07.14.664806

**Authors:** Grace A. Osborne, Lin Zhang, Feiyang Ma, Mehrnaz Gharaee-Kermani, Jessica Turnier, Amanda N. Victory, Amy Hurst, Bin Xu, Elisabeth A. Pedersen, Rachael Bogle, Celine Berthier, Vladimir Ognenovski, Mio Nakamura, Lam C. Tsoi, Allison C. Billi, Johann E. Gudjonsson, Benjamin Klein, Pei-Suen Tsou, J. Michelle Kahlenberg

**Author notes:** Corresponding author: J. Michelle Kahlenberg, MD, PhD,. These authors contributed equally (Grace A. Osborne, Lin Zhang).

## Abstract

Dermatomyositis (DM) is a rare yet devastating autoimmune disease characterized by inflammatory and vasculopathic changes in skin and muscle. DM and systemic lupus erythematosus (lupus) skin lesions have overlapping clinical and histopathological features, yet disparate responses to available therapeutics. DM skin disease is often relapsing and recalcitrant. To investigate DM immunopathogenesis, non-lesional skin, lesional skin, and circulating immune cells from DM patients were analyzed using single-cell RNA-sequencing. Samples were analyzed in parallel with lesional and non-lesional lupus skin, healthy control skin, and peripheral blood. We demonstrate a pervasive type I interferon (IFN) signature in DM stroma that persists in culture and is distinguished from lupus by upregulation of VEGF and IL-18 signaling in DM keratinocytes. Furthermore, endothelial cells (ECs) in lesional DM exhibit decreased proliferation that was not observed in lupus. Using cell communication networks, we identified a population of DM-specific monocytes interacting with non-proliferating DM ECs. Co-culture of monocytes from DM patients with ECs resulted in increased EC apoptosis inhibited by JAK1 blockade. JAK1 inhibition also resulted in reversal of DM-stromal and inflammatory signatures. Together, our data provide a comprehensive cross-disease characterization of lesional and non-lesional skin of DM compared to lupus and implicate monocyte-mediated EC dysfunction in DM vasculopathy and support JAK inhibition for refractory skin disease.

## INTRODUCTION

Dermatomyositis (DM) is a rare, chronic autoimmune disease that affects predominantly the skin and muscle but may also involve other organs, such as the lung. Skin disease is often relapsing and recalcitrant to therapy, even when systemic disease is well controlled(*1, 2*). Lack of knowledge regarding the pathogenesis of DM and drivers that instigate disease has delayed effective therapy development.

Like the skin of systemic and cutaneous lupus erythematosus (CLE) patients, IFNs are upregulated in lesional DM skin (*3–5*) and are also believed to play a pathogenic role in the destruction of muscle tissue (*6, 7*). Recent studies suggest myeloid-derived dendritic cells may be an important source for IFN-β in DM lesional skin (5); however, other cell populations have not been comprehensively studied. Moreover, our understanding of differences between DM and lupus skin lesions remains under-explored; yet, these differences are likely important to understand disease drivers and treatment responses.

Inflammation-associated vascular damage also contributes to the pathogenesis of DM. Loss of intramuscular microvessels accompanied by endothelial cell (EC) activation and poor regenerative capacity are hallmarks of DM-related endothelial cell dysfunction (*8, 9*). Early disease stages of DM have been associated with higher VEGF production by fascial endothelial cells (*10*), which may reflect ongoing vascular injury and attempts at regeneration (*11*). Additionally, circulating markers of endothelial cell injury are increased in DM patients and correlate with disease activity (*12, 13*). Thus, while endothelial dysfunction has been demonstrated in the muscle compartment, the role of ECs in the skin and the cellular mediators of activation and/or communication with immune cell populations have not been well-defined.

In this paper, we utilized single-cell RNA sequencing (scRNA-seq) to examine the cellular composition of paired lesional and non-lesional skin samples from DM patients with active skin disease and compared the findings to healthy control and non-lesional and lesional skin from lupus patients to comprehensively define the cellular makeup of the skin and to characterize mediators of inflammatory changes that contribute to disease. We also examined paired peripheral blood from the same DM patients to determine the origin of immune cell populations that contribute to EC activation and cell death. Overall, we found an IFN-rich stromal environment in DM that is distinct from lupus in upregulation of IL-18 and VEGF pathways. Further, DM skin exhibits EC senescence, and we demonstrate increased JAK-1-dependent EC death in DM that is likely driven by a DM-specific population of inflammatory monocytes. Our data provide detailed insights into DM pathogenesis and support a role for the use of JAK inhibition in refractory DM skin disease.

## RESULTS

### Comparison of epidermal and stromal cell populations from non-lesional and lesional skin of patients with dermatomyositis (DM) or lupus identifies disease-specific differences

To investigate the cellular composition and comprehensive transcriptional differences of lupus and DM skin, we performed scRNA-seq on lesional and sun-protected, non-lesional skin (upper thigh) of 11 lupus patients with active skin disease (5 with active discoid lupus erythematosus and 6 with active subacute cutaneous lupus erythematosus), and 8 DM patients with active skin disease (see **Fig 1a** for study design and **Supplemental Table 1** for patient details). Samples were analyzed in parallel with skin from 8 healthy controls (HC). The final dataset comprised 123,150 cells, with an average of 2,226 genes and 8,960 transcripts per cell. The cells were clustered based on differential expression of marker genes and visualized on a uniform manifold approximation and projection (UMAP) (**Fig 1b**). Cluster annotation was corroborated by overlapping the cluster markers with the canonical lineage specific genes reported in previous skin disease scRNA-seq studies (*14–18*) (**Fig S1a**). We recovered 11 major cell types, each found in DM (LDM), non-lesional DM (NDM), lesional lupus (LLE), non-lesional lupus (NLE), and HC (H) skin biopsies(**Fig 1c,d**). Cells in each type were derived from many samples across patient populations (**Fig S1b**).

**Figure 1.**
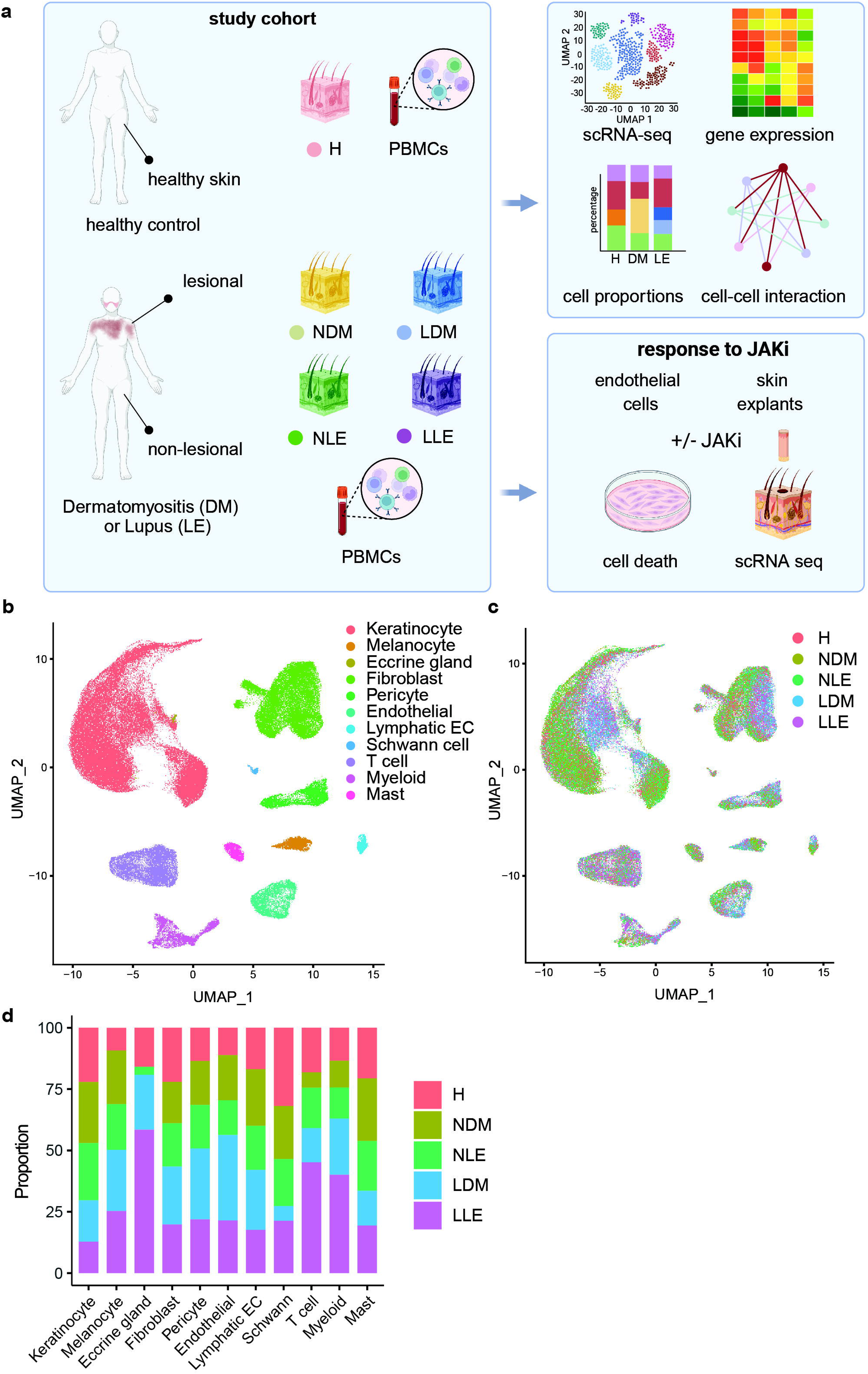
Single-cell RNA-sequencing (scRNA-seq) compares cell populations from non-lesional and lesional skin of patients with dermatomyositis (DM) to cutaneous lupus erythematosus (CLE) a. Schematic of human sample acquisition and experimental approaches throughout the study. b. UMAP plot of 123,150 cells colored by cell type c. UMAP plot of cells colored by disease state. H, HC skin. NDM, non-lesional dermatomyositis. NLE, non-lesional lupus. LDM, lesional dermatomyositis. LLE, lesional lupus d. Dot plot of representative marker genes for each cell type. Color scale, average marker gene expression. Dot size, percentage of cells expressing marker gene e. Bar plot of cell type proportions across disease states

Clustering by lesional skin state (i.e., lesional compared to non-lesional) was evident for many cell types, including keratinocytes (KC), fibroblasts (FBs) and ECs for DM and lupus lesions, suggesting similar transcriptional changes in these disease states (**Fig 1c,d**). Cell composition analysis revealed an increase in immune cell populations, including both T cell and myeloid lineages, in lesional lupus skin compared to LDM and HC (**Fig 1d**). There was also a notable increase in endothelial cells recovered in lesional DM skin compared to lesional lupus and HC (**Fig 1d**).

KCs constituted the majority of cells sequenced (61,008). Analysis of characteristic KC subtype markers identified 7 KC states: basal, spinous, supraspinous, granular, follicular, basal inflammatory, and spinous inflammatory (**Fig 2a,c**). Lesional skin from both lupus and DM dominated basal inflammatory and spinous inflammatory KC populations (**Fig 2b,c**). Consistent with previous work, basal inflammatory KC populations were also noted in non-lesional lupus KCs(*19*). This population was also present to a lesser extent in nonlesional DM skin (**Fig 2c**). To compare the transcriptomic differences in the basal inflammatory KC subpopulation by disease state, we performed differential expression analysis between lupus and DM non-lesional and lesional skin. When directly compared, lupus non-lesional basal inflammatory KCs exhibited some unique type I IFN-induced genes (*IFI27, IFI6*) whereas non-lesional DM basal inflammatory KCs exhibited unique expression of genes related to cell cycle, protein degradation and ribosomal function (*RPS26, UBE2S, EGR1*) (**Fig 2d**). Comparison of lesional basal inflammatory KCs identified a stress response signature (*S100A9, S100A8, S100A7*) with possible incorporation of KCs from the hair follicle (*K6b, K16*) in lupus lesions and heat shock proteins involved in mitochondrial signaling and stress response in DM (*DNAJA1, DNAJB1)* (**Fig 2e**). Total DEG analysis can be found in **Supplemental Tables 2 and 3**.

**Figure 2.**
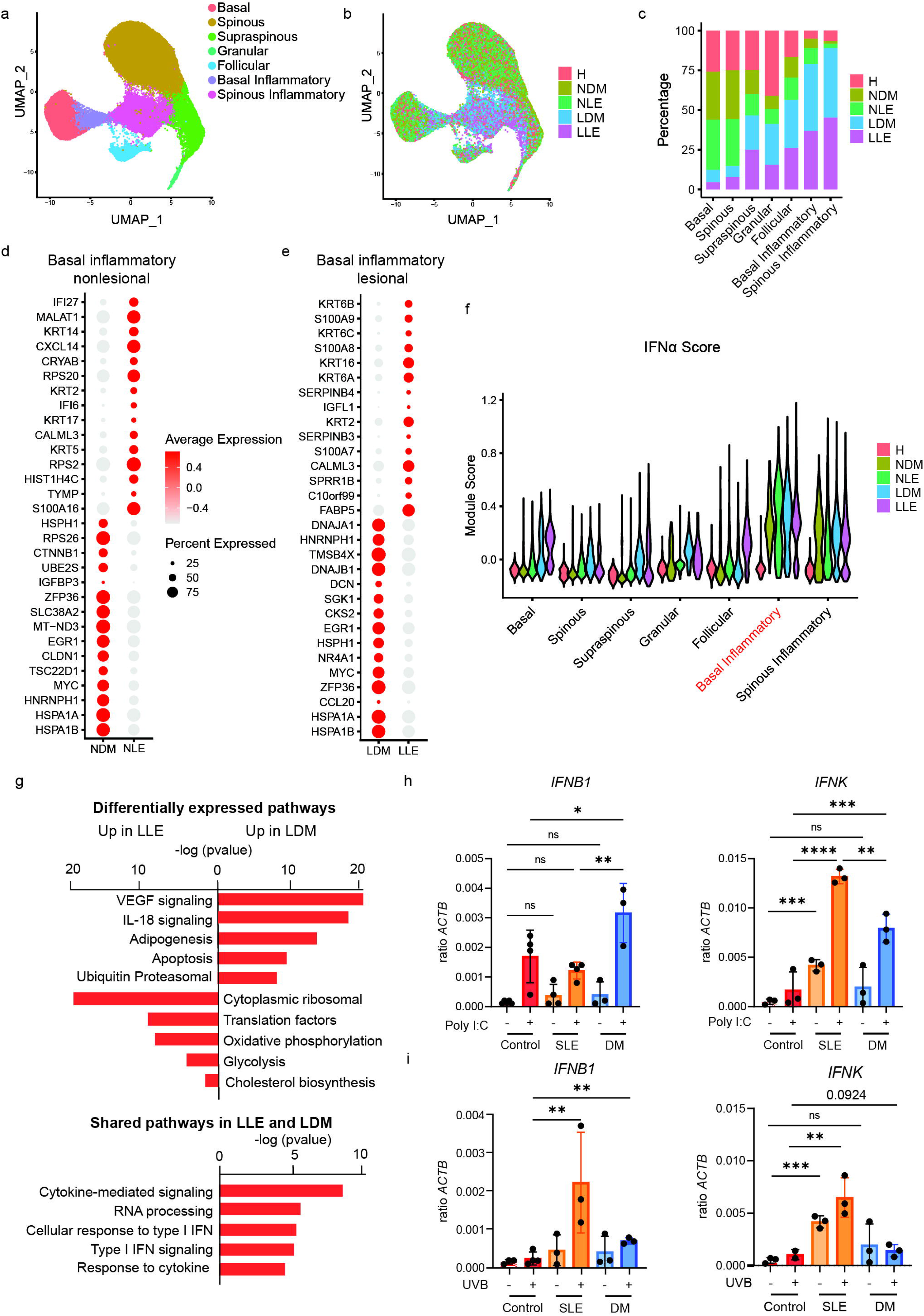
scRNA-seq analysis highlights differences in cytokine signature of DM and CLE stroma that persists in culture. a. UMAP plot of 61,008 keratinocytes (KCs) colored by sub-cluster b. UMAP plot of KCs colored by disease state c. Bar plot of number of KCs in each sub-cluster split by disease state d. Dot plot of the top differentially expressed genes (DEGs) upregulated and downregulated in basal inflammatory non-lesional disease states. Color scale, average marker gene expression. Dot size, percentage of cells expressing marker gene e. Dot plot of the top differentially expressed genes (DEGs) upregulated and downregulated in basal inflammatory lesional disease states. f. Violin plots of KC scores for the indicated cytokine modules split by disease state for each KC subcluster g. Bar graph of top upregulated pathways in LLE and LDM h-i. IFN gene expression by RT qPCR following stimulation of primary CTL, LUPUS, or DM (n=3 each) basal keratinocytes from skin samples with PolyI:C (10 ug/mL) (h) or UVB (50 mJ/cm^2^) (i), Ordinary one-way ANOVA followed by Šidák’s multiple comparison test or t test. Mean and SD. *P < 0.05, **P < 0.01, ***P < 0.001, and ****P < 0.0001.

We then wanted to compare cytokine exposures across diseases and lesional skin state to understand the relative dominance of inflammatory signals. The relatively shallow depth of scRNA-seq precludes direct examination of transcript levels for many cytokines implicated in DM and lupus – particularly type I IFNs. Thus, we calculated KC module scores derived from genes induced in cultured KCs upon stimulation with the indicated cytokines and generated violin plots displaying module scores for inflammatory cytokines relevant to skin and autoimmune diseases as we have previously reported (*19–22*) (**Supplemental Fig 2)**. While DM and LUPUS biopsies exhibited increases in cytokine responses in many KC clusters compared to control, the prominence of the type I IFN signature in basal inflammatory KCs for both disease states was particularly prominent with non-lesional lupus>lesional DM>lesional lupus>non-lesional DM >>healthy control (**Fig 2f**). Interestingly, follicular lesional lupus KCs also exhibited a high type I IFN signature, raising the question of whether IFN signaling plays a role in the pathogenesis of periadnexal inflammation and follicular plugging commonly noted in discoid lupus subtypes (**Fig 2f**). LDM exhibited relatively greater IL-1β in basal, supraspinous, follicular, and basal inflammatory subclusters compared to HC and lupus (**Supplemental Fig 2**).

To obtain a better overall understanding between lesional lupus and DM skin, we performed canonical pathway analysis using Ingenuity Pathway Analysis (IPA) software to compare the top 250 uniquely upregulated genes in the basal inflammatory KC cluster of LDM vs. LLE. Intriguingly, VEGFA-VEGFR2 signaling (p= 1.35×10^-19^) and IL-18 signaling (p=8.08×10^-17^) emerged as the most significantly enriched pathways in LDM DEGs whereas cytoplasmic ribosomal proteins, translation factors and mitochondrial oxidative phosphorylation and glycolytic metabolism pathways were enriched in lesional lupus basal keratinocyte DEGs (**Fig 2g**). Shared pathways in LLE and LDM using the top 250 genes upregulated in basal inflammatory keratinocytes in both diseases revealed enrichment of cytokine mediated signaling, RNA processing, cellular response to type I IFN, type I IFN signaling pathways, and response to cytokines (**Fig 2g**).

Given the limitations of scRNA-seq to detect type I IFNs, we then examined IFN production by primary basal keratinocytes isolated from non-lesional skin biopsies of patients with DM, lupus and HC. Consistent with our prior publications (*23, 24*), lupus KCs exhibited a significant chronic baseline upregulation of *IFNK* expression compared to HC KCs (**Fig 2h**). In contrast, there was not a significant increase in baseline *IFNK* or *IFNB* in DM KCs compared to HC KCs (**Fig 2h**). Stimulation with Poly(I:C) (10 ug/ml), which activates both TLR3 and RNA cytosolic sensing pathways induced both *IFNB* and *IFNK* in the lupus and DM KCs to a greater extent than HC. Intriguingly, a preference for *IFNK* was seen in lupus whereas *IFNB* production was more prominent in DM, reminiscent of other data acquired via tissue mass cytometry in DM immune populations(*3*). We then studied UV light exposure, another stimulus relevant to both DM and lupus as they are both photosensitive diseases. Following UVB stimulation, lupus KCs increased *IFNK* and *IFNB* production. DM KCs also increased transcription of these IFNs to a greater extent than HC but to a lesser extent than lupus (**Fig 2i**). Together, these data support type I IFN-driven epidermal changes in DM but to a lesser extent than those found in lupus patients.

We next analyzed FB populations. Annotation of these sub-clusters using published dermal FB marker genes (*25*) revealed SFRP2+, APOE+, TNN+, and CLDN1+ populations (**Supplemental Fig 3a-d).** The SFRP2+ cell population included two subpopulations SFRP2+PI16+ and SFRP2+COMP+ fibroblasts. SFRP2+PI16+ fibroblasts expressed high levels of *PI16* and *CCN5*, a repressor of TGFβ signaling (*26*). There were no notable differences between FB subclusters between disease states; however, a clear shift was noted in UMAP placement between non-lesional (comprising both LUPUS and DM) and lesional disease states (LUPUS and DM) (**Supplemental Fig 3b,c**). To determine the drivers between lesional and non-lesional disease states, we generated violin plots depicting FB cytokine module scores calculated using genes induced in cultured FBs stimulated by the indicated cytokines (**Supplemental Fig 3e**). Interestingly, both SRFP2+ FB populations and APOE+ FB from lesional DM were most uniquely distinguished by IFNα, IFNγ, and TNFα cytokine module scores (**Supplemental Fig 3e**). A list of DEGs between non-lesional and lesional SRFP2+ FB populations for DM and LUPUS samples are shown in **Supplementary Tables 4-7**. These data indicate that inflammatory fibroblast education occurs in both DM and LUPUS and that lesional FBs may be more affected in DM vs. LUPUS skin.

### Endothelial cells are activated and demonstrate increased cellular senescence in lesional DM compared to other disease states

While both LUPUS and DM patients exhibit endothelial dysfunction(*9, 27*), an important clinical feature of many DM patients is nailfold capillary changes (*9, 28*), an indication that endothelial cells may play a unique role in DM disease pathogenesis. We thus compared transcriptome changes seen in endothelial cells between HC, LUPUS, and DM. Sub-clustering analysis of 5,611 dermal blood vascular endothelial cell (EC) identified 11 sub-clusters with distinct marker genes (**Fig 3a-d**). We identified three EC subtypes based on transcriptional signatures: arteriole (*GJA4, SEMA3G*), venule (*SELP*) and capillary (*RGCC*) (**Fig 3b-d**). EC0 appears as a transitional capillary->venule cluster and thus is of interest as this is the site of immune cell migration and interaction. LDM ECs predominated in three sub-clusters of ECs: EC0, EC8, and EC10 (**Fig 3e**). Interestingly, these EC subtypes showed evidence of activation (*SELE* and *ICAM1*), upregulation of IFN-induced genes (*ISG15*) and inflammatory regulators (*TNFAIP3* (encodes A20)) genes (**Fig 3d**).

**Figure 3.**
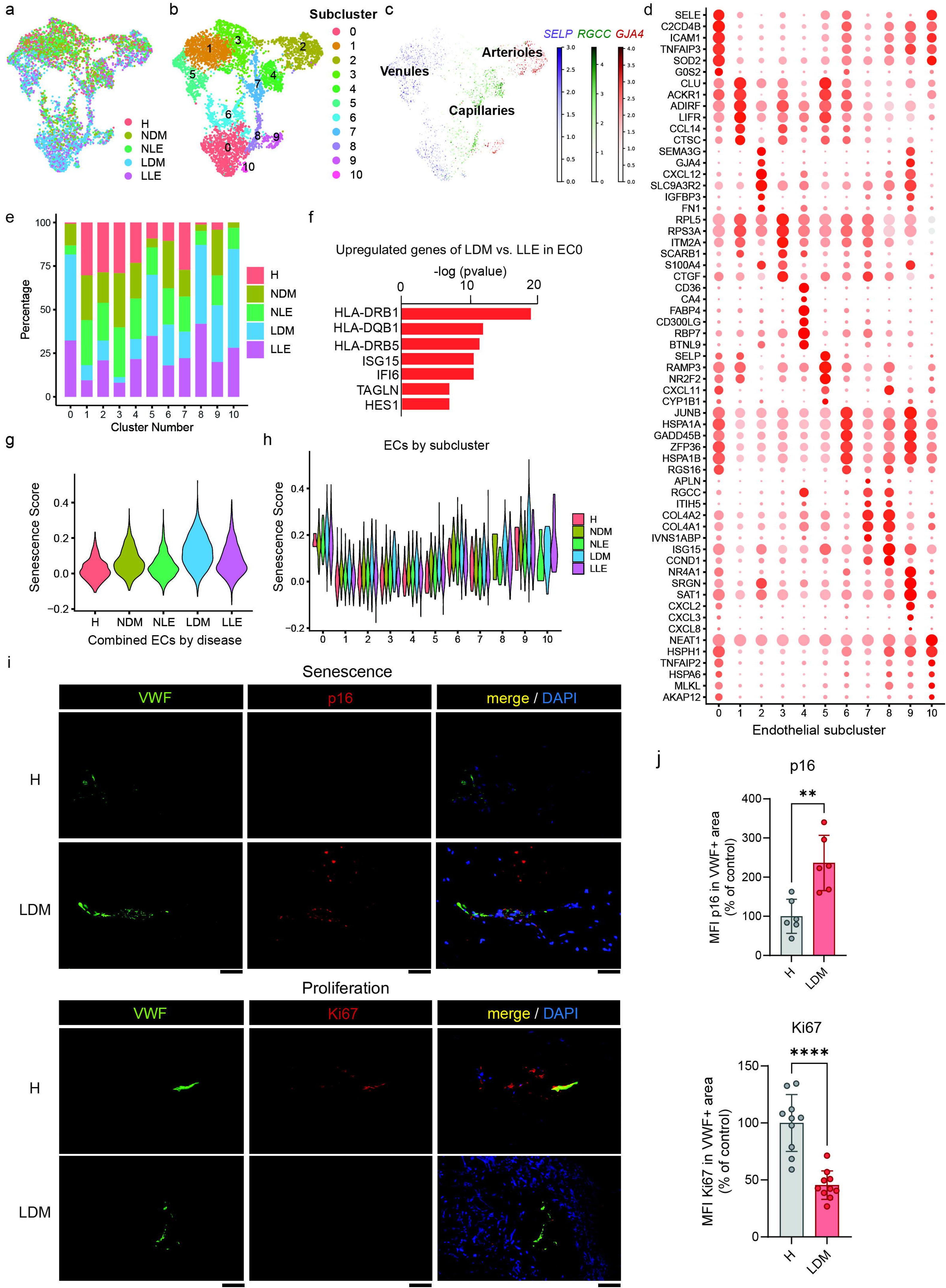
Endothelial cells are activated and demonstrate increased cellular senescence in lesional DM compared to other disease states. a. UMAP of endothelial cells (ECs) colored by disease state b. UMAP of ECs colored by sub-cluster c. schematic of endothelial cell classification and representative gene marker d. Dot plot of the representative marker genes for each subcluster. Color scale, average marker gene expression. Dot size, percentage of cells expressing marker gene e. Bar plot of number of ECs in each sub-cluster split by disease state f. Bar plot of upregulated genes of lesional DM vs. lesional LE in endothelial subcluster 0 g. Violin plot of senescence score of combined endothelial cell by disease state h. Violin plot of senescence score of endothelial cells by subcluster i. Representative immunofluorescence of frozen HC and lesional DM skin biopsies (n=3). j. Quantification of immunofluorescence for p16 and Ki67 using mean fluorescence intensity (MFI) across vWF+ cells, dots represent average of 3 different areas per vessel. Student’s t test, **P < 0.01 and ****P < 0.0001.

Given that both DM and LUPUS exhibited expansion of capillary cell clusters, we performed canonical pathway analysis via IPA of the top 200 upregulated genes in lesional DM ECs compared to lesional lupus ECs in cluster EC0. In lesional DM, crosstalk with myeloid cells was emphasized as IL-10 signaling, antigen presentation, and macrophage activation were the top 3 significant pathways compared with lesional lupus (**Supplementary Fig 4a**). When comparing the transcriptional differences of this subcluster in LDM compared to LLE, the most differentially expressed genes were those of the major histocompatibility complex II (MHC) *HLA-DRB1* (p=7.8×10^-17^), *HLA-DQB1* (p=3.34×10^-8^), and *HLA-DRB5* (p=1.22×10^-8^), (**Fig 3f**). IFN response genes *ISG15* and *IFI6* were also upregulated in the EC0 subcluster of LDM compared to LLE, as were *HES1,* a gene involved in NOTCH signaling, and *TAGLN,* a gene involved in fibrosis (**Fig 3f**). IPA was used to identify upstream regulators of lesional DM versus lupus transcriptional changes in EC0. Interestingly, HSF1, a molecule that has been reported to drive endothelial prostaglandin production (*29*) and anticoagulation factors in endothelial cells (*30*), and SATB1, a regulator of chromatin remodeling during active cell death (*31*) were significant upstream regulators in DM, whereas TNF signaling was a more dominant upstream driver of changes unique EC0 cells from lupus samples (**Supplementary Fig 4b**).

Senescent phenotypes have been identified to play a strong role in vasculopathy in other autoimmune diseases, such as scleroderma (*32*). Intriguingly, increased oxidative stress in the capillary ECs was suggested by the upregulation of superoxide dismutase (*SOD*) in EC0 (**Fig 3d**). We thus applied a senescence score generated from a list of 79 genes annotated as associated with cellular senescence based on published work (*33–36*) or cellular senescence annotation in the Reactome database to the EC clusters. The full gene list for generation of the senescence score can be found in Shi *et al* (*33*) and includes senescence-associated secretory phenotype genes and senescent gene families. Remarkably, the non-lesional and lesional DM ECs exhibited increased senescence scores when compared to non-lesional and lesional lupus ECs, respectively (**Fig 3g**). The senescent signature was highest in EC0 and EC9 (**Fig 3h**). To examine what may be driving these changes, we used IPA to identify the top signals predicted to serve as upstream regulators for the genes induced in lesional ECs taken as a whole. Consistent with known inducers of endothelial dysfunction and senescent phenotypes, LDM samples had much higher predictive values of signaling by IFNγ, IL1β, NONO (a regulator of transcription and RNA splicing), IL27, type I IFNs and TNF compared to LLE (**Supplementary Fig 4c**). These data suggest that in lesional DM, inflammatory signaling contributes to differential phenotypes in DM vs. lupus cutaneous ECs, including a propensity towards senescence that could lead to vasculopathy.

To further examine the senescent phenotype at the protein level, we performed immunofluorescence staining on skin sections derived from three HC and three lesional DM skin biopsies for p16 (marker of senescence) and Ki67 (marker of proliferation). Von Willebrand Factor (VWF) was used to stain endothelial cells. Consistent with our transcriptional data, we identified increased EC costaining with the senescence marker p16 and a decrease in costaining with the proliferation marker Ki67 in lesional DM EC compared to HC (**Fig 3j).**

### T peripheral helper cells are expanded in lesional DM compared to lupus and HC

We next examined the immune cells in the skin. Sub-clustering and annotation based on established marker genes identified 7 T cell subsets: CD4 T cell (CD4T), T regulatory cell (Treg), T peripheral/follicular helper (Tph/fh), CD8 T cell (CD8T), Tissue resident memory (Trm), Natural killer cell (NKC) and Innate lymphoid cell (ILC) (**Fig 4a-c**). CD4T cells were the predominant T cell subset, comprising 42-60% for all disease states followed by CD8 T cells which were13-14% of all T cells (**Fig 4 d-e**). Cells of one sub-cluster were distinguished by expression of ISGs and were therefore designated IFN T cells (**Fig. 4c**). An increased percentage of all IFN T cells derived from LDM compared to LLE. IFN T cells were also recovered from non-lesional lupus, but not non-lesional DM skin (**Fig 4d-e**).

**Figure 4.**
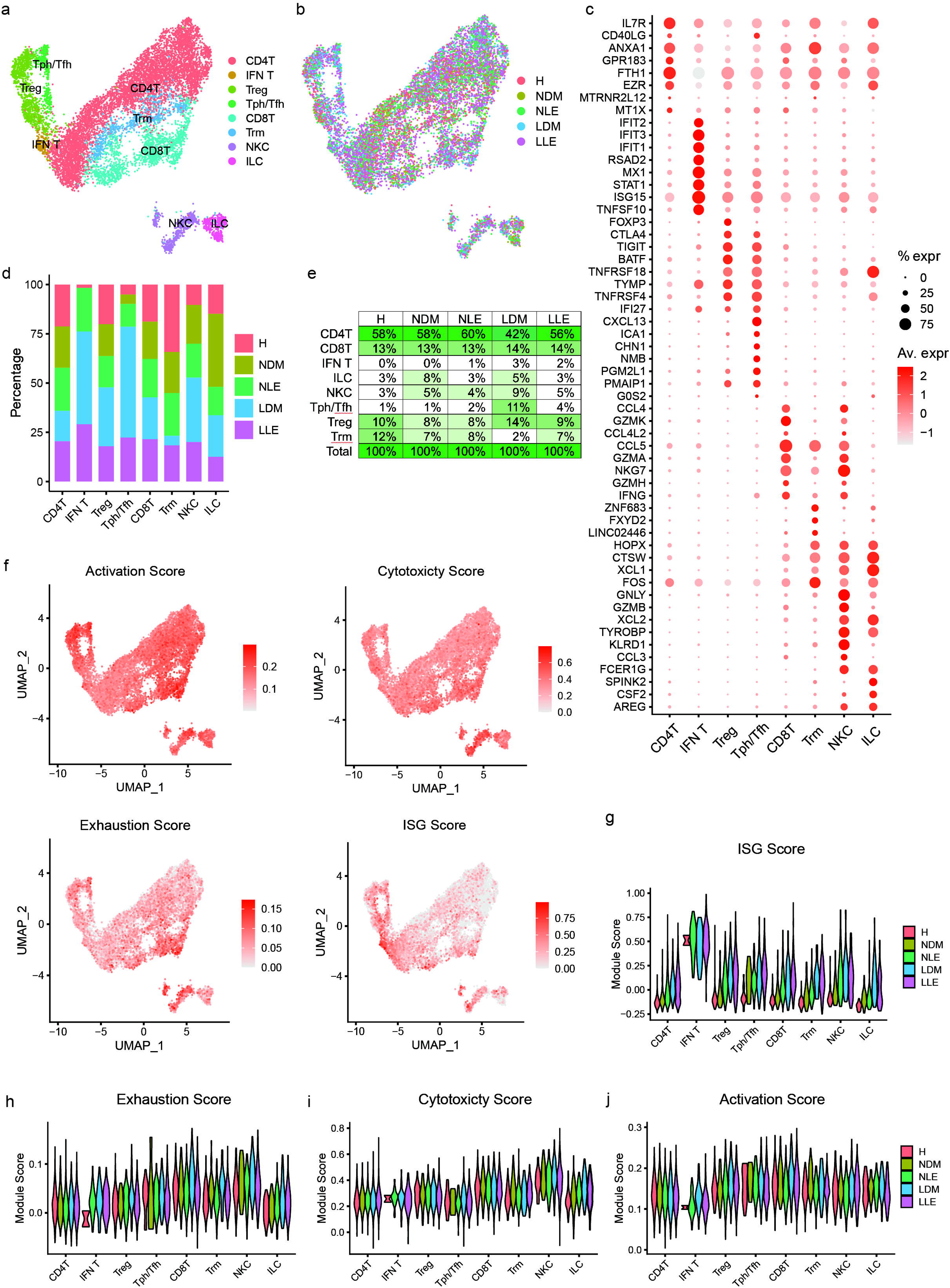
T peripheral helper cells are expanded in lesional DM compared to lupus and healthy control. a. UMAP of T cells from skin biopsies by T cell subtype b. UMAP of T cells from skin biopsies by disease state c. Dot plot of representative marker genes for each T cell subtype d. Bar graph of number of T cells in each subtype by disease state e. percentage of T cell type by disease state f. UMAP of activation, cytotoxicity, exhaustion and ISG scores g-j. Violin plots of ISG (g), exhaustion (h), cytotoxicity (i) and activation (j) scores in T cell subsets by disease state.

The Tph subcluster was notably expanded in lesional DM patients compared with other disease states. The abundance of these cells varied by disease state at 1% for HC, 1% for NDM, 11% for LDM, 2% for NLE, and 4% for LLE T cells (**Fig 4e**). This subcluster was distinguished by expression of *CXCL13*, encoding a B cell-attracting chemokine that has been implicated in the loss of immune tolerance of B-cells in idiopathic inflammatory myopathies (reviewed in Zanframundo et al. (*37*)). To investigate the characteristics of the DM-predominant Tph subcluster, we generated features plots displaying the module scores for T cell characteristics (activation score, cytotoxicity score, exhaustion score and interferon-stimulated gene (ISG) score). The Tph subset was predominantly activated without strong cytotoxic or exhaustion scores. CD8 T cells were activated as well but demonstrated increased exhaustion and cytotoxicity scores. Cytotoxicity scores were highest in NK cells, but exhaustion scores were also elevated (**Fig 4f**). ISG scores were enhanced in IFN T cells, confirming this phenotype (**Fig 4f**). Comparing across diseases, ISG scores were elevated in all immune cell populations in DM and LUPUS lesions, with non-lesional LE exhibiting scores higher than non-lesional DM in NK cells, Tregs, and CD4 T cells (**Fig 4g**). Evaluation of exhaustion scores identified a trend for more pronounced exhaustion in lesional DM CD8T, Trm cells, and NK cells (**Fig 4h**). Cytotoxicity and exhaustion scores did not vary much between diseases (**Figure 4i, j**). In summary, lesional DM skin has an increase in Tph populations that are transcriptionally similar to lesional lupus regarding activation. Cytotoxic populations seem to be functionally similar in DM vs. lupus lesions with possibly greater exhaustion in cytotoxic cells such as CD8T and NK cells in lesional DM.

### The Langerhans cell population is expanded and inflammatory in nonlesional DM skin

We then evaluated myeloid cells, the other major immune cell type detected in our skin samples. Sub-clustering and annotation of myeloid cells from skin biopsies revealed 11 myeloid cell subsets and a B cell subset with largely distinct marker genes (**Figure 5a-c).** Subsets included Langerhans cells (LC; *CD207*, *CD1A*), classical type 1 dendritic cells (cDC1; *DNASE1L3*, *C1orf54*), classical type 2 dendritic cell subset A (cDC2A; *CCL17*, *CCL19, BIRC3*) and subset B (cDC2B; *CLEC10A*, *IL1B*), CD16+DCs, interferon dendritic cells (IFN DC), lipid-associated macrophages (LAM; *APOE*, *APOC1*), plasmacytoid dendritic cell-like (pDC-like; *PPP1R14A*), and plasmacytoid dendritic cells (pDCs; *GZMB*, *JCHAIN*). B cells were primarily derived from lesional lupus skin, consistent with the literature (*38, 39*). Myeloid cell subsets showed variability in representation among disease states. Among the most striking differences was that LC were captured primarily from DM samples (**Fig 5d)**, with 52% and 10% of all LC deriving from non-lesional and lesional DM samples, respectively (**Fig 5e**). Consistent with a known reduction of LCs in lupus skin (*40*), only 2% of LCs derived lesional lupus samples, whereas non-lesional lupus skin was similar to HC (16% vs 14%) (**Fig 5e**). Immunofluorescence staining for CD207 in frozen non-lesional DM skin confirmed that LCs were increased compared to HC and NLE (**Fig 5f**). In NDM LCs, there were 139 significant upregulated genes (Log_2_FC > 0.25, FDR adjusted p_value < 0.05), compared to HC LCs. Predicted pathways included IL-1 signaling, TCR signaling, antigen processing and presentation (**Fig 5g**, **Supplemental Table 8**). 156 significant upregulated genes (Log_2_FC > 0.25, FDR adjusted p_value < 0.05) were found in LDM LCs vs HC LCs (**Supplemental Table 9**), and IPA analysis suggested the top significant signaling pathways were related to type I IFN (**Fig 5g**). These data indicate that LC phenotype may be more consistent with monocyte-derived inflammatory LCs in DM patients (*41*).

**Figure 5.**
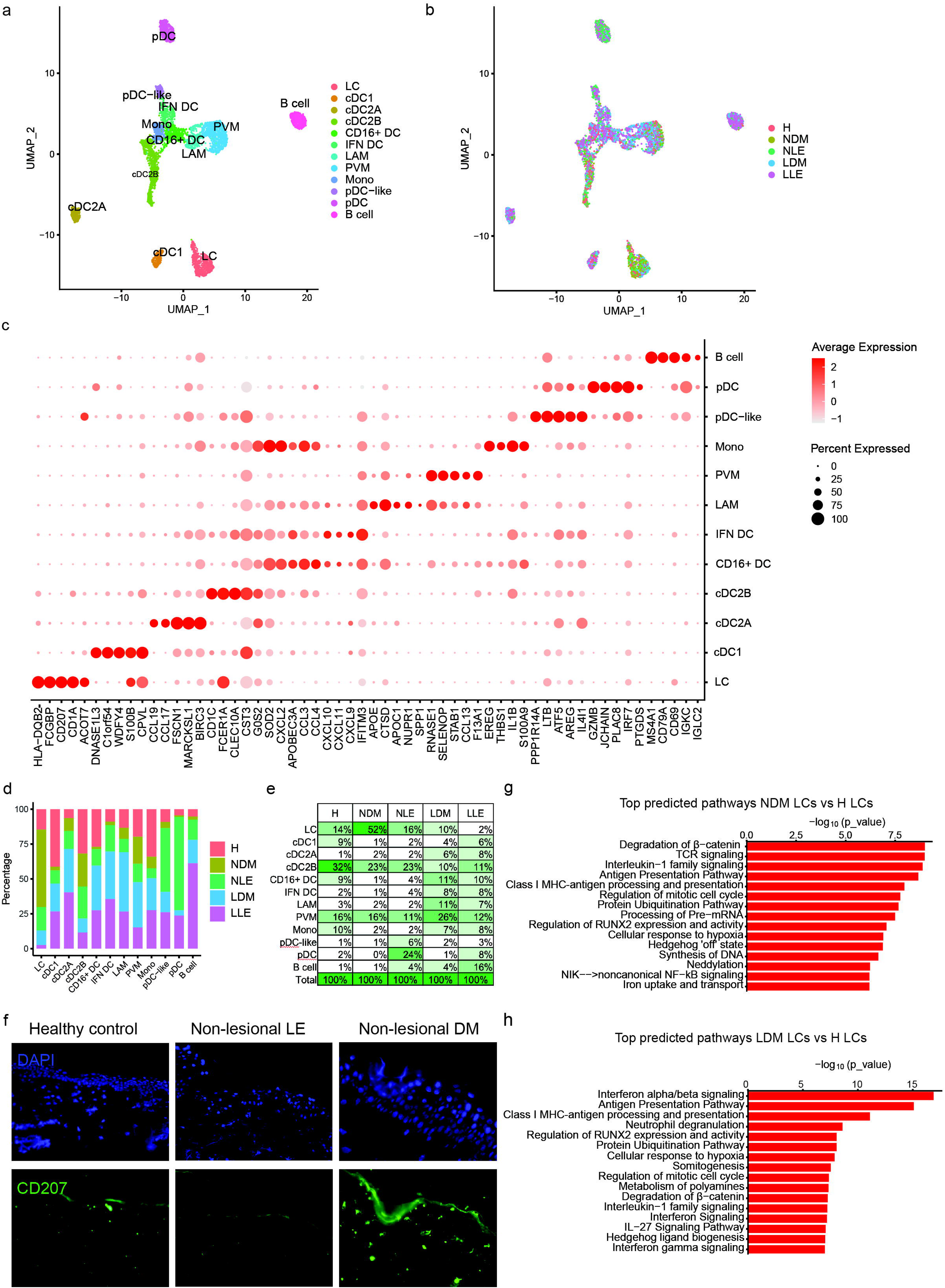
Major differences in myeloid cell subsets are detected in lesional and non-lesional skin of DM compared to lupus. a. UMAP of skin derived myeloid cells by subtype b. UMAP of skin derived myeloid cells by disease state c. Dot plot of representative marker genes for each myeloid subtype d. Bar graph of myeloid cells by subtype e. Percentage of myeloid cells by disease state f. Representative immunofluorescence of Langerhans cell marker CD207 in frozen DM lesional skin biopsies (n=3) g. Bar graph indicating activation Z-scores of predicted upregulated pathways in NDM vs HC Langerhans cells (LC). h. Bar graph indicating activation Z-scores of predicted upregulated pathways in LDM vs HC Langerhans cells (LC).

### Peripheral Blood of DM patients exhibits a unique, inflammatory classical monocyte population

After analysis of the skin clusters, we then examined peripheral blood mononuclear cells (PBMCs) by scRNA-seq from five of the same DM patients and all LUPUS patients as well as all HC (**Fig 6a,b, Supplemental Fig 5a)**. Interestingly, a monocyte subcluster from DM PBMCs (DMP) showed a distinct shift on the UMAP away from HC and LUPUS peripheral blood (HP and LEP) (**Fig 6a,b and Supplemental Fig 5b**). To understand the transcriptional changes that accompanied this shift, monocytes were then selected for further sub-clustering. We identified 6 monocyte subsets with distinct marker genes (**Fig 6c-e**). Intermediate and classical monocytes were in clusters 1-4 and nonclassical monocytes in cluster 5 (high expression of *FCGR3A*). DM patient samples constituted the vast majority of cells in the DM-specific cluster Mono0 (**Fig 6f**). Numerous DM patients contributed to this monocyte population, both treatment naïve and treated (**Supplementary Table 1, Supplementary Fig 5c**). Overall, we identified Mono0 as a subset of classical monocytes as it expresses very high levels of CD14 (**Supplementary Figure 5d**). Cluster markers identified expression of numerous inflammatory cytokines and chemokines in Mono0, including *CXCL1* and *CXCL5*, which are unusual in HC monocyte populations, as well as expression of *FGF2*, an important activator of ECs and neovascularization (*42*)(**Fig 6e)**. Critically, using IPA, upstream cytokine regulators driving gene expression within Mono0 vs. other monocyte clusters were identified to include inflammatory cytokines such as IL-1β, TNFα, IL17A, IL-6, and IL-18 (**Supplementary Figure 5e**), suggesting circulating monocytes in DM patients are educated strongly by inflammatory conditions-even more strongly than circulating LUPUS monocytes. To further compare this, we examined the top canonical pathways enriched in DM Mono0 compared to other monocyte subclusters and identified coronavirus (IFN) signaling, IL-10 pathways (composed of *CCL2, CCL20, CCL3, CCL4, CXCL1,* and *IL6* as notable genes in the pathway), and mitochondrial dysfunction as skewed pathways in Mono0 (**Figure 6G**). Relationships between PBMC and skin myeloid cell data were then further explored via aggregation of circulating and skin-associated myeloid cells. (**Fig 6h).** Sub-clustering of the aggregated myeloid cells revealed that Mono0 monocytes were found amongst the skin populations only in DM patients (**Fig 6h, red arrowhead**), suggesting that these cells are not only in circulation but also interface with the skin.

**Figure 6.**
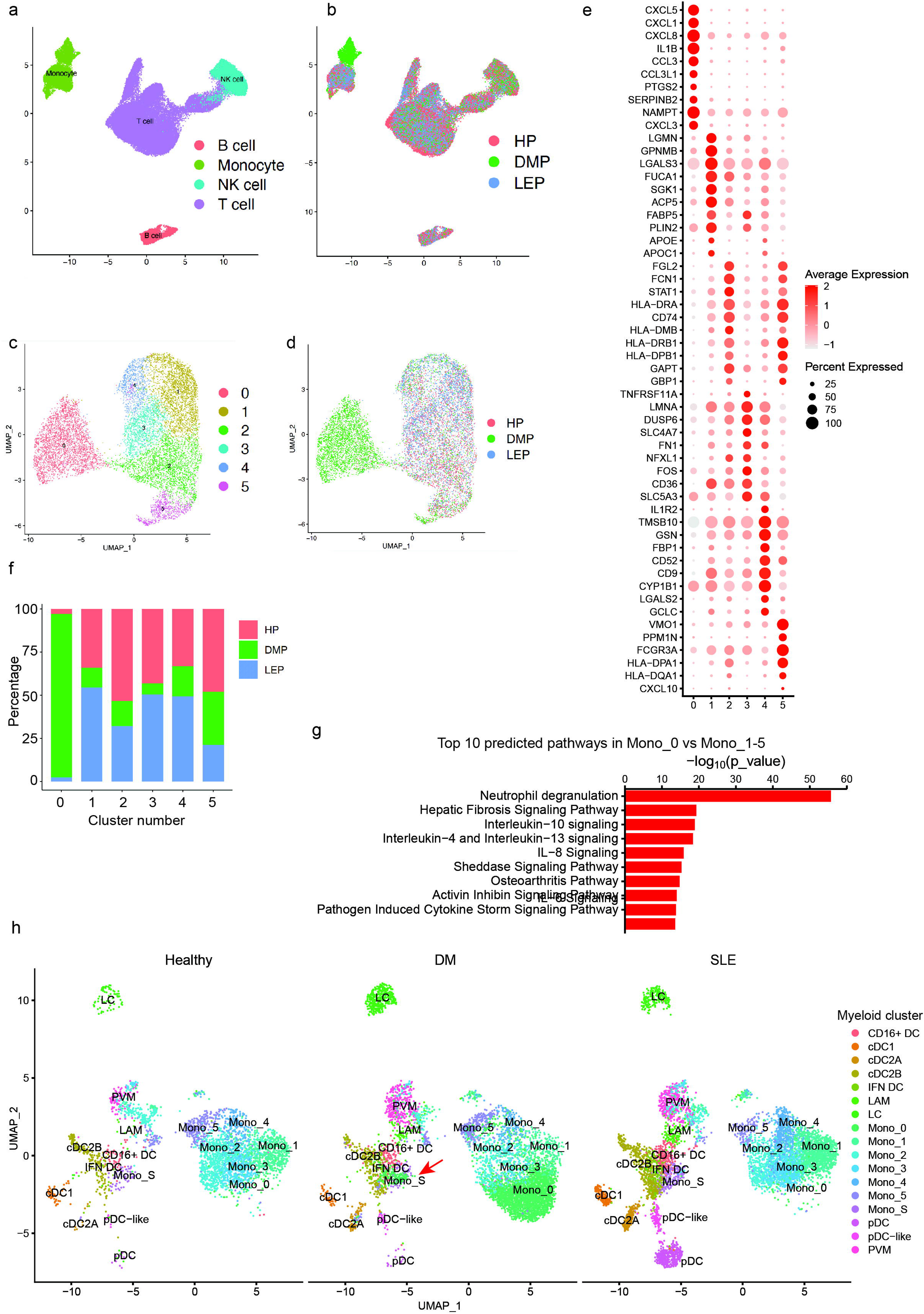
Peripheral blood monocytes migrate into skin of lesional DM. a. UMAP of PBMC derived myeloid and B cells by cell type b. UMAP of PBMC derived myeloid and B cells by disease state c. Bar graph of PBMC derived myeloid cells by subtype d. Dot plot of representative marker genes for each myeloid subtype e. UMAP of PBMC monocytes by subtype f. UMAP of PBMC derived monocytes by disease state g. Dot plot of representative marker genes by monocyte subtype h. Bar graph of monocyte subtype by disease state i. Bar graph of upregulated pathways in DM specific monocyte subpopulation 0 j. UMAP of combined skin and PBMC derived myeloid cells by cell subtype k. UMAP of combined skin and PBMC derived myeloid cells by disease state

### Ligand-receptor analysis demonstrates DM-enriched cell-cell interactions among peripheral monocytes and dermal ECs

As Mono0 cells are found in DM skin, we then wanted to examine which cell populations they were interacting with. We thus used CellChat to examine cellular crosstalk. When acting as senders (expressing ligands), Mono0 showed prominent interactions with ECs (**Figure 7A**); when acting as receivers (expressing receptors), Mono0 showed strong interactions with fibroblasts (**Figure 7b**). The strong mono0/EC interactions were of interest given the inflammatory and senescent changes in DM ECs (EC0,6,8,9,10) (**Figure 3**). We used Cellphone DB to specify critical ligand-receptor interactions between Mono0 and ECs that were upregulated in LDM vs. HC skin. Crosstalk was noted to be robust and was notable for Mono0 production of VEGFA and B, TNFα, IL-6, IL-1β, TNFSF14 (LIGHT) and TNFSF12 (TWEAK) (**Supplementary Figure 6**). Given that capillary changes including dropout are prominent feature of DM nailfolds (*28*) and musculature (*43*), it was intriguing to note the Mono0 production of inflammatory mediators, such as IL-6 and TNF family members, known to promote EC apoptosis and senescence (*44*). We thus tested whether monocytes from DM patients exhibited an enhanced ability to kill ECs. We co-cultured DM and HC CD14+ monocyte populations isolated via negative selection from PBMCs (n=3 each) and compared their effects on ECs using a dye for activated caspase3/7. When compared to age and sex-matched HC monocytes, DM monocytes induced EC death at a significantly higher rate than HC, suggesting that DM monocytes may contribute to vascular dysfunction and inflammation in DM skin and possibly other organs (**Figure 7c**).

**Figure 7.**
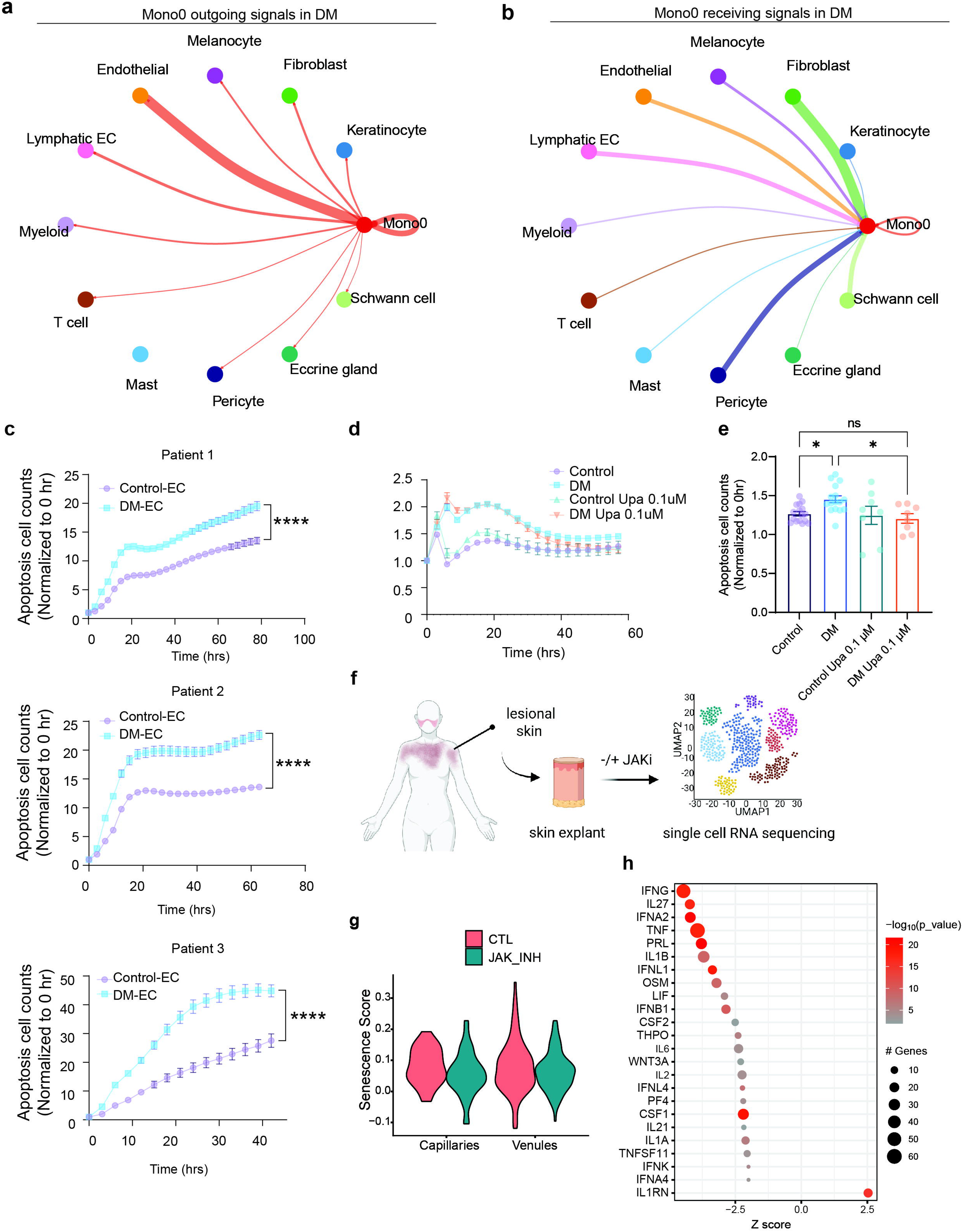
Peripheral monocytes from dermatomyositis patients result in increased apoptosis of dermal blood vascular endothelial cells. a. Receptor ligand analysis showing circulating ncMo ligand to EC receptors b. Receptor ligand analysis showing circulating ncMo receptor and EC ligands c. Plot of EC apoptosis in monocyte killing assay d. Plot of EC apoptosis in monocyte killing assay with inhibition of JAKSTAT signaling with 0.1 uM upadacinitib (Upa). e. Quantification of (d). f. Schematic of lesional skin explant study: Explants were treated with JAKi and processed for scRNAseq g. Senescence scores in untreated (CTL) and JAKi-treated (JAK_INH) capillaries and venules. *p<0.05 h. Dot plot of the top 25 upstream regulators enriched among DEGs in untreated DM versus JAKi-treated DM. Color scale, −log_10_(*P* value) from the enrichment analysis. Dot size, number of DEGs corresponding to each upstream regulator; negative *z* score, enriched in CTL; positive, enriched in JAKi-treated.

Various newer medications have been proposed for treatment of dermatomyositis, including JAK inhibitors, which are useful for refractory skin disease (*45*) and are known to block many cytokines including IL-6 (*46*). We thus tested the JAK1 inhibitor upadacitinib in our culture system and found that it significantly attenuated the increased apoptosis of ECs (**Figure 7 c-e**). A TWEAK inhibitor did not have beneficial effects on monocyte-mediated cell killing (data not shown). We then took a broader look at the effects of upadacitinib on the skin as a whole. We studied explanted lesional skin from two untreated patients with active DM and treated the biopsies overnight with or without upadacitinib followed by scRNA-seq (**Figure 7f**). Cell quality was good and cell populations were similar to our initial UMAP (**Supplemental Figure 7A**). Importantly, with upadacitinib treatment, *AKAP12*+ ECs exhibited a reduction in capillary senescence scores (**Figure 7g, Supplemental Figure 7B,C**). This was accompanied by reduced inflammatory signaling as defined by inhibition of IFNγ, type I IFN, TNF, and IL6 signaling in upadacitinib-treated samples (**Figure 7h**). In contrast, no reduction in monocyte inflammatory pathways was seen with upadacitinib treatment (**Supplemental Table 10**). Thus, endothelial dysfunction is partly reversed via JAK1 inhibition, potentially through inhibition of monocyte interactions such as IL-6 rather than direct inhibition of monocytes themselves.

## METHODS

### Human sample acquisition

For skin biopsies for single cell RNA-seq, 8 DM patients with active skin disease (**Supplemental Table 1**) were recruited for this study, all of whom contributed lesional and non-lesional (sun-protected skin of the buttock) 6mm punch skin biopsies. A diagnosis of DM was confirmed for all patients via the 2017 European League Against Rheumatism/American College of Rheumatology classification criteria(*47*). 11 patients with active CLE +/- lupus (10 with associated systemic lupus and 1 CLE only) were included for comparison. 8 HC were also recruited for skin biopsy. For endothelial cell-death assays, additional DM and age and sex-matched HC subjects were recruited for blood draws only. An additional two active DM patients were recruited for explant studies with upadacitinib. The study was approved by the University of Michigan Institutional Review Board (IRB), and all patients underwent written, informed consent. The study was conducted according to the Declaration of Helsinki Principles.

### Single-cell RNA library preparation, sequencing, and alignment

Generation of single-cell suspensions for scRNA-seq was performed as follows: Skin biopsies were incubated overnight in 0.4% dispase (Life Technologies) in Hank’s Balanced Saline Solution (Gibco) at 4°C. Epidermis and dermis were separated. Epidermis was digested in 0.25% Trypsin-EDTA (Gibco) with 10U/mL DNase I (Thermo Scientific) for 1 hour at 37°C, quenched with FBS (Atlanta Biologicals), and strained through a 40μM mesh. Dermis was minced, digested in 0.2% Collagenase II (Life Technologies) and 0.2% Collagenase V (Sigma) in plain medium for 1.5 hours at 37°C, and strained through a 40μM mesh. Epidermal and dermal cells were combined in 1:1 ratio, and libraries were constructed by the University of Michigan Advanced Genomics Core on the 10x Chromium system with chemistry v3. Libraries were then sequenced on the Illumina NovaSeq 6000 sequencer to generate 150 bp paired end reads. Data processing including quality control, read alignment (hg38), and gene quantification was conducted using the 10x Cell Ranger software. The samples were then merged into a single expression matrix using the CellRanger aggr pipeline.

### Cell clustering and cell type annotation

The R package Seurat (v4.0.3)(*48*) was used to cluster the cells in the merged matrix. Cells with less than 500 transcripts or 100 genes or more than 10% of mitochondrial expression were first filtered out as low-quality cells. The NormalizeData function was used to normalize the expression level for each cell with default parameters. The FindVariableFeatures function was used to select variable genes with default parameters. The ScaleData function was used to scale and center the counts in the dataset. Principal component analysis (PCA) was performed on the variable genes, and the first 30 PCs were used in the RunHarmony function from the Harmony package(*48, 49*) to remove potential batch effect among samples processed in different libraries. Uniform Manifold Approximation and Projection (UMAP) dimensional reduction was performed using the RunUMAP function. The clusters were obtained using the FindNeighbors and FindClusters functions with the resolution set to 0.6. The cluster marker genes were found using the FindAllMarkers function. The cell types were annotated by overlapping the cluster markers with the canonical cell type signature genes. To calculate the disease composition based on cell type, the number of cells for each cell type from each disease condition were counted. The counts were then divided by the total number of cells for each disease condition and scaled to 100 percent for each cell type. For clustering of combined myeloid cells (skin myeloids and PBMC monocytes), the R package Seurat (v4.3.0.1)(*48*) was used to re-cluster the combined myeloid cells (skin myeloids and PBMC monocytes). The six subsets from the PBMC monocytes (Fig 6C) were annotated as Mono_0 – Mono_5 within the combined myeloid UMAP (Fig 6H). The monocytes within the skin myeloids (Fig 5C) were labelled Mono_S within the combined myeloid UMAP (Fig 6H).

### Module score analysis

Module scores for FBs and KCs (IFN_α_ score, IFN_γ_ score, IL-1_β_ score, TNF_α_ score), ECs (senescence score) and T cells (activation score, cytotoxicity score, exhaustion score, and IFN-stimulated gene score) were calculated using the AddModuleScore function on the gene signatures previously published as follows: For FBs and KCs, the gene signatures are available within the supplementary data files s3 and s2, respectively, in Billi *et al*. (19). For ECs, the full gene list for generation of the senescence score can be found in Shi *et al.* (*33*). For T cells, the activation, cytotoxicity, exhaustion, and IFN-stimulated gene signatures are available within the supplemental table 3 in Dunlap *et al*(*50*).

### Cell type sub-clustering

Sub-clustering was performed on the abundant cell types. The same functions described above were used to obtain the sub-clusters. Sub-clusters that were defined exclusively by mitochondrial gene expression, indicating low quality, were removed from further analysis. The subtypes were annotated by overlapping the marker genes for the sub-clusters with the canonical subtype signature genes.

### Cell-Cell communication analyses

CellChat (v2.1.2)(*51*) was used to find out the interactions between the cluster 0 of PBMC monocytes (Mono_0) with the different types of skin cells. CellPhoneDB (v2.0.0)(*52*) was used for Ligand-Receptor analysis. The cells were separated by their disease classifications (H, LDM, NDM, LLE, NLE). Pairs with *P* value > 0.05 were filtered out from further analysis. To compare among the five disease conditions, each pair was assigned to the condition in which it showed the highest interaction score. The number of interactions for each cell type pair was then calculated. In supplementary Fig 6, the top interactions from Mono_0 (Ligands) to skin AKAP12+ ECs (EC0, EC6, EC8, EC9, EC10) were plotted in Circos plots using the R package “circlize.”

### Cell Culture

Primary human keratinocytes were established from non-lesional lupus and DM patients and healthy control punch biopsies as previously described (15). Cells were maintained in culture with Epilife medium and were passaged at 50% confluency to avoid differentiation prior to study. For functional assessment, cells were treated with 10 ug/mL poly (I:C) or were irradiated with 50mJ/cm^2^ UVB (310nm) via a UV-2 irradiator (Tyler Industries, Alberta, Canada). Cells were incubated in fresh medium for 6 hrs or 24 hrs, respectively, followed by harvest and RNA isolation for real-time PCR (*IFNB1* FW 5’3’: ACGCCGCATTGACCATCTAT RV 5’3’: GTCTCATTCCAGCCAGTGCT, *IFNK* FW 5’3’: GTGGCTTGAGATCCTTATGGGT, RV 5’3’: CAGATTTTGCCAGGTGACTCTT).

### Immunofluorescence

Immunofluorescence was performed on frozen sections obtained from skin biopsies from HC or non-sun exposed, non-lesional lupus or non-lesional DM patient skin. Thawed sections were fixed in −20 **°**C acetone, washed, and blocked with 5% BSA for 1 hr at RT. Slides were placed in primary antibody overnight at 4°C, washed and placed in secondary antibody for 1.5 hrs at RT. Antibodies used CD207 (Biolegend clone 929F3.01). Images were acquired using Zeiss Axioskop 2 microscope and analyzed by SPOT software. Images are representative of n=3 control and n=3 LUPUS, n=3 DM.

### Monocyte isolation and killing assay

Human dermal microvascular endothelial cells (Lonza) were seeded in 96-well plates (Corning) at a density of 2 × 10^4^ cells per well and grown overnight. On the day of the assay, monocytes (40,000 cells/well) were isolated from HC and DM patients using StemCell negative selection kits (StemCell technologies, #19359) following Ficoll separation of PBMCs. Monocytes were added to each well of endothelial cells together with caspase-3/7 reagent (1:2000 dilution, Essen Bioscience). Endothelial cells were imaged at 10-fold magnification in an IncuCyte S3 Live Cell Analysis System (Sartorius) at 37°C with 5% CO_2_. Images were acquired every 3 hours, 4 images per well. Data were analyzed using IncuCyte analysis software to detect and quantify the number of green (apoptotic) cells per image. Data were plotted using GraphPad Prism software. Data was presented as mean ± SEM, statistical analysis was done using two-way ANOVA.

### Statistical analysis and data sharing

Statistical analysis was performed in GraphPad for functional assays using 2-way Anova or Student’s t-test where indicated, where p-values less than 0.05 were considered significant. As for the scRNA-seq data, we performed, using the FindMarkers function in the Seurat package, differential gene expression analyses between H and LDM, H and NDM, or LLE and LDM, where an FDR-adjusted p-value less than 0.05 was considered statistically significant. Ingenuity pathway analysis was applied to the differentially expressed genes to determine the canonical pathways and the potential upstream regulators, and those with a Z-score of ≥ 2 or ≤ −2 were considered significant. Single cell RNA-seq data from DM and lupus not already shared on GEO (accession number GSE186476) will be shared following publication of the manuscript.

## DISCUSSION

In this work, we have provided a comprehensive understanding of the cellular similarities and differences between DM and lupus non-lesional and lesional skin using scRNA-seq and identified several important unique attributes in DM patients. These include dysfunctional states of keratinocytes, an increase in inflammatory LCs, and a unique, proinflammatory circulating monocyte population that may contribute to a propensity for endothelial cell death and senescence in the tissue.

We have identified that while type I IFN signaling is a relevant pathway in both diseases, KC IFN signaling may be less prominent in DM than in CLE skin. Interestingly, DM keratinocytes exhibited a propensity for IFN-β responses over IFN-κ, which has also been observed in other cell types in previous studies(*3, 53*). Notably, IFN-β correlates with cutaneous disease activity in DM(*54*). The reasons for a preference for IFN-β vs. other type I IFNs in DM could include epigenetic changes, genetic regulation or other yet undefined mechanism and should be further investigated. Importantly, trials using antibodies targeting IFN-β in DM have completed early phase studies, with promising results (*55*).

In addition to type I IFN, DM keratinocytes show enhanced IL-18 signaling, in line with previous findings linking DM and an IL-18 gene signatures using microarray data of lesional skin(*5*). IL-18 represents a pleiotropic cytokine and its expression in epithelial cells is promoted by multiple upstream signals, including type I and II IFN(*56*). Secretion of IL-18 is driven by inflammasome activation, which has not been studied in DM skin disease, but IL-18, IL1β and NLRP3 expression levels in muscle tissue have been reported as elevated in DM(*57*). Canonical downstream effects of IL-18 are MyD88-dependent, leading to activation of MAPK and NF-κB signaling(*56*). Unlike IL-1β, IL-18 can be easily detected in peripheral blood and it will be interesting to dissect the role of IL-18 signaling in DM disease overall.

Other cell populations in DM skin also exhibited some differences from CLE. Both lupus and DM have a pronounced inflammatory fibroblast component; however, SRFP2+PI16+, APOE+, and SRFP2+COMP+fibroblasts from lesional DM exhibit robust cytokine signatures. Our analysis suggests they are signaling strongly to monocytes as they enter the tissue, and they may be playing an important role in inflammatory education of immune cells in the skin. Whether these fibroblasts are contributing to future development of calcinosis or tissue atrophy (all potential complications or consequences of DM skin lesions), requires further study.

Vasculopathy represents a life-threatening manifestation in DM, and clinically apparent nail fold capillary changes occur in up to 89% percent of DM patients (*58*). Our evaluation of ECs identified pronounced inflammatory changes in DM capillary EC clusters driven by type I IFN and HSF1 signaling whereas LE ECs were dominated by TNF signaling. This suggests that DM capillaries have different sources and consequences of inflammatory signals compared to LE, and these differences may be reflected by clinical features and treatment responses. While endothelial dysfunction has been demonstrated in circulating endothelial progenitors and in DM-associated myositis(*9, 59*), our results identify pronounced senescence and apoptosis of DM skin endothelial cells, which contrasts with the lupus phenotype in our data. Multiple cytokines and other external stressors promote senescence, including IL-6, TNF and type I IFNs(*60, 61*). In DM, we identify type I IFNs and IL-6 as potential drivers. In addition, the Mono0 population exhibits activation of multiple inflammatory pathways including IL-18. Indeed, IL-18 was shown to contribute to endothelial cell progenitor dysfunction in lupus and DM(*9*). Whether monocyte-derived IL-18 is a contributor to vascular dysfunction in DM skin needs to be further explored.

Intriguingly, the most prominent difference in cell populations was noted in circulating PBMCs in which a population of hyperinflammatory CD14+ monocytes was noted in DM (“cluster 0” monocytes). These cells exhibited strong expression of inflammatory cytokines and chemokines, were found to infiltrate DM lesional skin, and had strong interactions with endothelial cells. This contrasts with what has been previously reported in lupus skin, where CD16+ DCs played a more influential role(*19*). Our data thus confirms and builds on the role for CD14+ monocytes in DM skin. Indeed, tissue mass cytometry has found that CD14+ monocytes in DM skin correlate with cutaneous disease severity(*3*). Another study in juvenile DM found mitochondrial dysfunction in CD14+ peripheral blood monocytes contributing to type I IFN signaling(*62*), which was also a prominent signaling in our adult DM CD14+ Mono0 cluster. Our data further suggest that this population of highly inflammatory CD14+ monocytes contributes to endothelial dysfunction in DM, which can be targeted through JAKi. This is in line with a previous report showing that JAK1/2 inhibition using ruxolitinib restored type I IFN-mediated vascular network disruption in DM(*63*). Previous trials showed clinical efficacy of JAKi in DM(*45*), but cellular mechanistic analyses of tissues has been lacking. Our results showed JAK-dependent, but TWEAK-independent killing of endothelial cells by patient-derived monocytes, corroborating the importance of therapeutically targeting monocyte-derived signals in DM. Furthermore, JAKi treatment reduced the senescent phenotype of endothelial cells in DM skin explants.

Together, our results provide single cell resolution of DM skin and functional evidence that targeting monocyte-EC interaction via JAK inhibition represents a new avenue to treat vasculopathy and likely simultaneously improve other features of disease. Future clinical trials will dissect which patients benefit from JAKi and whether early intervention might prevent endothelial dysfunction in DM.

### Study Funding

Funding for this work was received through the Lupus Research Alliance (to JMK), Bristol Myers Squibb (to JMK and JEG), and the National Institutes of Health NIAMS via R01 AR071384 and K24 AR076975 (to JMK), AI130025 (to JEG), the U-M Skin Biology and Diseases Resource Center P30 AR075043 (to JEG) and NIAID via P01 AI179251 (to JMK and JEG) and R01AI183620 (to JEG and PT). Funding was also received through the Taubman Institute Innovative Program (to JMK and JEG), the Department of Defense (to PT), and the LEO Foundation (to PT). JLT was supported by a NIAMS K23 Career Development Grant (K23AR080789). BK was supported by the German Research Foundation (KL3612/1-1)

## Acknowledgements

We are grateful to the patients of the Michigan Lupus Program and our dermatomyositis patients for participating in this study and donating precious skin to help us understand the biology of their diseases. All single cell RNA-sequencing was performed by the U-M Advanced Genomics Core and we are grateful for their assistance. We also acknowledge Kelsey McNeely for her technical assistance with CellChat.

## Supplementary Tables

Table S1. Dermatomyositis patient demographics

Table S2. Lesional DM vs. lesional lupus DEG analysis in basal inflammatory KCs, |log2FC| is 0.1.

Table S3. Non-lesional DM vs. non-lesional lupus DEG analysis in basal inflammatory KCs |log2FC| is 0.1.

Table S4. Non-lesional DM vs. non-lesional lupus DEG in all SRFP2+ FB, |log2FC| is 0.1.

Table S5. Non-lesional DM vs non-lesional lupus DEG in SRFP2+PI16+ FB, |log2FC| is 0.1.

Table S6. Non-lesional lupus vs. lesional lupus DEGs in all SRFP2+ FB, |log2FC| is 0.1.

Table S7. Non-lesional lupus vs. lesional lupus DEGs in SRFP2+PI16+ FB, |log2FC| is 0.1.

Table S8. Non-lesional DM vs. healthy DEGs in Langerhans cells, |log2FC| is 0.1.

Table S9. Lesional DM vs. healthy DEGs in Langerhans cells, |log2FC| is 0.1.

Table S10. Downregulated pathways in skin monocytes after JAKi treatment.

**Supplementary Fig 1.**

○ a. Dot plot of representative marker genes for each cell type. Color scale, average marker gene expression. Dot size, percentage of cells expressing marker gene
○ b. UMAP plot of cells colored by disease state.

**Supplementary Fig 2.**

○ a. UMAP plot of fibroblasts (FB) colored by sub-type
○ b UMAP plot of FB colored by disease state
○ c. Bar plot of number of FBs in each sub-type split by disease state
○ d. dot plot of marker gene expression by FB subtype
○ e. cytokine plots by FB subtype

**Supplementary Fig 3.**

○ a. Upstream regulators of lesional DM vs. lesional lupus in endothelial cell subcluster 0

**Supplementary Fig 4.**

○ a. Significantly enriched pathways in lesional DM vs. lesional lupus ECs
○ b. Comparison of upstream regulators in lupus and DM EC0 clusters
○ c. Comparison of upstream regulators in DM vs. lupus ECs.

**Supplementary Fig 5.**

○ a. Cluster markers for PBMCs from DM, lupus, and HC subjects
○ b. Proportions of cell populations from PBMCs from DM, lupus, and HC subjects
○ c. UMAP of monocyte clusters by donor
○ d. Expression of indicated genes across monocyte clusters
○ e. Upstream regulators of mono0 vs. other monocyte subclusters

**Supplementary Fig 6.**

○ Interactions between DM mono0 subcluster and ECs as defined as ligand from mono0 and receptor in ECs in DM skin.

**Supplementary Fig 7.**

○ UMAP of cell clusters from skin explants
○ Cluster markers from skin explants
○ UMAP of endothelial cells from skin explants
○ *AKAP12* expression across EC clusters.

