## Supplemental Table 1 for "Cross-disease comparison of dermatomyositis and lupus skin identifies inflammatory monocytes and JAK-1 signaling as drivers of vasculopathy in dermatomyositis"

| **Age (y)** | **Sex** | **Race and ethnicity** | **Positive Antibodies** | **Disease Manifestations at time of Biopsy** | **Autoimmune medications at time of biopsy** | **Disease duration at time of bx** |
| --- | --- | --- | --- | --- | --- | --- |
| **DM patients for scRNA seq** | | | | | |  |
| 30 | F | White, non-Hispanic | Anti-MDA5, Anti SSa, Anti Ku (weakly positive) | myopathy, gottrons papules, photosensitivity, periungual erythema | IVIG, HCQ | 10 years |
| 59 | F | White, non-Hispanic | Anti-PL7 | photosensitivity, myopathy | Rituximab, cellcept, HCQ, IVIG | 3 years |
| 31 | F | White, Hispanic | ANA 1:640, speckled | myopathy, dysphagia, wong-varient rash | cellcept, IVIG | 6 years |
| 61 | F | Other | anti-TIF-1γ, anti-Ssa/Ro52, ANA >1:2,560 speckled | myopathy, rash, nail bed changes, alopecia | none | 11 years |
| 32 | F | White, non-Hispanic | ANA 1:640 speckled, AntiTIF1γ | rash , myopathy | HCQ, prednisone 7.5 mg | 2 years |
| 65 | F | White, non-Hispanic | ANA 1:320 homogeneous | rash, myopathy | rituxan | 13 years |
| 29 | F | White, non-Hispanic | ANA 1:640, speckled, Anti-RNP, Anti-cN-1A | rash, amyopathic | none | 3 weeks |
| 73 | F | White, non-Hispanic | ANA 1:1280, speckled | rash, amyopathic | plaquneil, Ivig, pred 5mg daily | 6 years |
| 66 | F | White, non-Hispanic | ANA 1:2560 speckled, +SSA/Ro-52,TIF-1γ | rash, dysphagia, myopathy | Rituxan, MTX, IVIg, plaquenil | 18 months |
| **SLE/CLE patients for scRNA-seq** | | | | | |  |
| 42 | F | Black, non-Hispanic | ANA, dsDNA, RNP, SSA/Ro, SSB/La, Ribo-P | Skin rash (DLE), ITP | None | 15 years |
| 42 | F | Black, non-Hispanic | ANA, dsDNA, Smith, RNP, SSA/Ro, SSB/La, cardiolipin IgM and IgG, β2GP1 IgM and IgA | Skin rash (DLE) | ASA, belimumab IV monthly, MMF, prednisone 5 mg daily | 20 years |
| 48 | F | White, Hispanic | ANA, dsDNA, Smith, RNP, SmRNP, SSA/Ro, Ribo-P, Chromatin | Skin rash (SCLE) | Dapsone 50 mg daily, HCQ 200 mg BID | 30 years |
| 48 | F | White, non-Hispanic | SSA/Ro, β2GP1 IgM | Skin rash, arthritis (SCLE) | HCQ, prednisone 7.5 mg daily, quinacrine, tofacitinib | 4 years |
| 60 | F | White, non-Hispanic | ANA, SSA/Ro | Skin rash (DLE) | none | 5 years |
| 64 | M | White, non-Hispanic | none | Skin rash (SCLE) | Apixaban, prednisone 5 mg daily, quinacrine | 3 years |
| 65 | F | White, non-Hispanic | ANA, dsDNA, β2GP1 IgA | Oral ulcers, Raynaud’s, APS, skin rash (ACLE/SCLE) | HCQ, warfarin | 1 year |
| 20 | F | Asian | ANA 1:320 homogeneous, Smith, dsDNA | Skin rash (DLE), alopecia, arthritis | HCQ | 4 years |
| 34 | F | White, non-Hispanic | ANA >1:2560, homogenous, dsDNA, chromatin, SSA/Ro | Skin rash (DLE), sicca, arthritis | AZA, prednisone 4 mg daily, belimumab | 2 years |
| 57 | F | White, non-Hispanic | ANA 1:640, speckled, SSA/Ro | Skin rash (SCLE), sicca, arthritis | MMF, HCQ | 3 years |
| 74 | F | White, non-Hispanic | ANA >1:2560, SSA/Ro pattern, SSA/Ro | Skin rash (SCLE) | HCQ | 1 year |
| 44 | F | White, non-Hispanic | ANA, SSA/Ro, dsDNA | Skin rash (DLE), arthritis, alopecia, thrombocytopenia, serositis | HCQ, prednisone 5 mg daily | 23 years |

**Table S1. Patients used for single cell RNA seq studies.** ANA, antinuclear antibody; APS, antiphospholipid syndrome; ASA, aspirin; BID, *bis in die* (twice daily); DLE, discoid lupus erythematosus; DM, dermatomyositis; dsDNA, double-stranded DNA; HCQ, hydroxychloroquine; ITP, immune thrombocytopenia; MMF, mycophenolate mofetile; PO, *per os* (by mouth); RNP, ribonucleoprotein; SCLE, subacute cutaneous lupus erythematosus; SLE, systemic lupus erythematosus.
